# Evolutionary history determines population spread rate in a stochastic, rather than in a deterministic way

**DOI:** 10.1101/2020.07.16.206268

**Authors:** Mortier Frederik, Masier Stefano, Bonte Dries

## Abstract

Fragmentation of natural landscapes results in habitat and connectedness loss, making it one of the most impactful avenues of anthropogenic environmental degradation. Populations living in a fragmented landscape can adapt to this context, as witnessed in changing dispersal strategies, levels of local adaptation and changing life-history traits. This evolution, however, can have ecological consequences beyond a fragmented range. Since invasive dynamics are driven by the same traits affected by fragmentation, the question arises whether fragmented populations evolve to be successful invaders.

In this study we assess population spread during three generations of two-spotted spider mite (*Tetranychus urticae*) population in a replicated experiment. Experimental populations evolved independently in replicated experimental metapopulations differing only in the level of habitat connectedness as determined by the inter-patch distance.

We find that habitat connectedness did not meaningfully explain variation in population spread rate. Rather, variation within experimental populations that shared the same level of connectedness during evolution was larger than the one across these levels. Therefore, we conclude that experimental populations evolved different population spread capacities as a result of their specific evolutionary background independent but of the connectedness of the landscape. While population spread capacities may be strongly affected by aspects of a population’s evolutionary history, predicting it from identifiable aspects of the evolutionary history may be hard to achieve.

## Introduction

Movement is integral to the life of all organisms and a principle driver of species distributions, spread and eventually the dynamics of ecosystems (Jeltsch et al., 2013). Environmental change and habitat loss put a heavy pressure on population persistence. One way to manage this pressure is to move to other locations with more suitable and benign environmental conditions (O’Connor, Selig, Pinsky, & Altermatt, 2012; Parmesan, 2006). Effective conservation policy requires knowledge on how fast and how likely a particular population can keep up with a changing landscape. These insights can similarly inform agricultural pest and infectious disease management. Many organisms are also deliberately or accidentally introduced outside their ancestral range. They sometimes manage to establish and spread further (Renault, Laparie, McCauley, & Bonte, 2018). In the past, this has spawned a series of invasive species that replaced their native counterparts (Mckinney & Lockwood, 1999). Predicting species invasion risk has therefore become a major theme in invasion biology.

The predictability of evolutionary change or ecological dynamics has historically been rather poor (Pigliucci, 2002). As such, predictability in population spread has gathered some interest but has been strongly debated as well (Giometto, Rinaldo, Carrara, & Altermatt, 2014; Melbourne & Hastings, 2009). Central to an accurate forecasting is the availability of reliable predictors. Population spread is affected by characteristics of the landscape but also by traits that determine movement and population growth (Angert et al., 2011; Fisher, 1937). Movement will determine how efficiently the landscape can be crossed while other life-history traits will determine the build-up of populations and eventually the number of the potentially spreading individuals. Spread itself induces a non-random distribution of these traits within the range that as a result accelerates spread. Dispersive phenotypes accumulate at the edge through spatial sorting and more reproductive phenotypes are selected for at the range’s edge by a process termed spatial selection (Burton, Phillips, & Travis, 2010; Fronhofer & Altermatt, 2015; Shine, Brown, & Phillips, 2011; Szücs et al., 2017). Whereas selection can act on the evolution of these traits, they are equally conditional non-adaptive processes such as genetic drift or linkage disequilibrium with adaptive traits. Moreover, a population’s historical context greatly influences its ecology in the present (Maris et al., 2018). Selection pressures and other evolutionary forces of past environments shaped the current traits of each population. The population’s historical environmental background is therefore expected to leave a signature on the population spread dynamics that may be predictable to a certain extent.

An important feature of the environment affecting the evolution of demography and movement is its overall level of habitat fragmentation (Cheptou, Hargreaves, Bonte, & Jacquemyn, 2017). Fragmentation usually is the direct result of habitat loss. But habitat fragmentation results in further stresses on natural populations, one of them being the increasing distance between patches of habitat. We will call this connectedness henceforth. Populations living in these increasingly less connected habitat patches will experience elevated dispersal costs (Bonte et al. 2012). As a consequence, less dispersal is expected to evolve, which leads to a decreased connectivity as expressed by a decreased amount of successful dispersers between spatially separated patches (Tischendorf & Lenore, 2001). Because changes in connectivity directly feedback with changes in local densities (Cheptou et al., 2017), growth rates and stress resistance can evolve as well (De Roissart, Wang, & Bonte, 2015; Bonte et al. 2018). While selection should lead to convergence in traits among populations experiencing the same spatial context, other factors may lead to more stochasticity in trait changes and the emerging population dynamics. First, connectedness loss predominantly coincides with a decrease in patch size. The resulting smaller populations experience an increased genetic drift and can lead to the loss of adaptive traits. Second, lower connectivity directly decreases gene flow among populations, leading to a direct loss of genetic variation (Lenormand, 2002) and an increased genetic load within populations (Ingvarsson, 2001).

Based on the above, we could expect populations inhabiting strongly connected benign landscapes to spread overall faster relative to those from less connected ones because of their higher dispersal abilities. On the other hand, evolution of stress-related traits may substantially lower the costs of dispersal in the less connected landscapes (Bonte et al., 2012; Cheptou et al., 2017). This may lead to an equal or even faster population spread for populations that share an evolutionary history in the poorly connected landscapes. Independently of the exact direction and magnitude of these effects, we hypothesize that population spread should be predictable in relation to the spreading population’s evolutionary history.

Eco-evolutionary dynamics predominantly show how the dynamic interplay between trait evolution and ecological dynamics in the same environment (Hendry, 2016). Quantifying the impact of trait-changes in one kind of environment on the dynamics in another environment are key to invasion theory (e.g. Bonte & Bafort 2018), but to date virtually unknown from an empirical or natural perspective. We therefore quantified dispersal propensity and reproductive rate prior to the experiments to compare how informative this trait perspective is compared to the evolutionary background of populations. Building on a long-term experimental evolution experiment (Masier & Bonte 2020), we quantified population spread dynamics of two-spotted spider mite (*Tetranychus urticae*) populations for 2-3 generation, thereby simulating the take-off of an invasion. By using replicated mesocosms that experienced the same or another level connectedness, as well as replicated range spread tests for each of these experimental mesocosms, we are able to quantify the predictability of early population spread (Giometto et al., 2014; Melbourne & Hastings, 2009), and thereby to estimate the importance of evolution for spread dynamics in a new environment. Overall, our results show that evolution affects population spread rate to a sizable extent but that the historical level of habitat fragmentation is an unconvincing predictor.

## Methods

### Experimental system

We tested population spread in *Tetranychus urticae* Koch (two-spotted spider mite) populations. The species is a cosmopolitan phytophagous herbivore known from >900 plant species (Navajas et al., 2002). This species is used as a model in ecology and evolutionary biology. The rapid population growth, ease of maintaining populations in a lab and the known genomics (Grbić et al., 2011) are all advantages for performing such research. For this experiment, we used an in-house lab population which had been used in other experiments (Alzate, Bisschop, Etienne, & Bonte, 2017; Bisschop, Mortier, Etienne, & Bonte, 2019; De Roissart, Wang, & Bonte, 2015; Van Petegem et al., 2018)

We maintained mites on *Phaseolus vulgaris* L. Prelude (bean) plants and leaf patches at all times. Bean is an optimal host for the spider mites, with little in the way of defense. Mites are never found to perform better on other hosts compared to bean, even when the mites locally adapted to that host for a prolonged period (Alzate et al., 2017). We created optimal resource conditions in the evolutionary and population spread setups for dynamics to not be affected by resource maladaptation.

### Evolutionary history

We evolved mites in lab-controlled mesocosms in a metapopulation spatial composition for 18 months. Mesocosms differed in the interpatch distance. The replicated mesocosms are described in more detail in Masier et al. (2019). In short, each evolutionary arena consisted of a 3×3 grid of bean leaf patches (5cmx5cm) that were connected by parafilm® bridges of 0.5cm wide to all adjacent patches (fig. 1). Horizontal and vertical bridges were all 4 cm, 8 cm or 16 cm long, determining the connectedness treatment of the mesocosm. The distance between bean patches mostly determined the dispersal mortality risk of a mite moving between patches. Each inter-patch distance treatment was replicated five times. During the 18 months of experimental evolution, leaf patches in each mesocosm were refreshed weekly. (Masier & Bonte, 2019) reported the evolution of the same dispersal propensity in the different connectedness treatments. However, the more connected mesocosms evolved a later dispersal timing and a greater starvation resistance.

**Figure 1:**
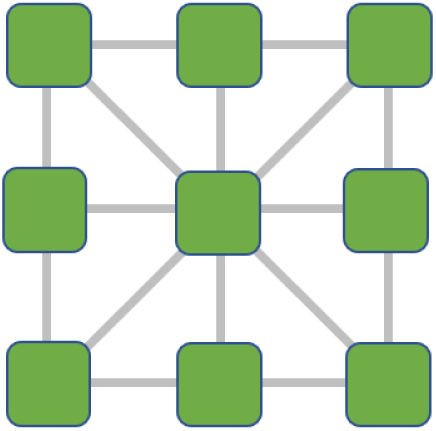
spatial configuration of the mesocosm landscape

After 18 months, we transferred 45 mature females to a bean leaf from each mesocosm: five from each local patch. In the few cases of low local population sizes, less than five mites were sampled to not compromise the viability of that local population. This bean leaf with transferred mites rested on wet cotton wool in a petri dish (150mm diameter) aligned with paper towel strips (30°C, 16:8h L:D photoperiod). We let these mature females lay eggs for 24h to form a synchronized next generation to perform the dispersal propensity and reproductive success tests with. Afterwards, all females from a mesocosm were transferred to a bean plant to breed a large enough number of individuals in the next generation for the population spread assessments. Both the leaf in the petri dish and bean plant provided a common garden for the mites used in their respective tests in order to control for maternal effects and effects of developmental plasticity.

### Population spread

We sampled 40 individuals from the common garden plant of each mesocosm and placed them in a population spread arena. We replicated this three time to have three independent population spread assessments per evolved mesocosm. Some of the whole plants used as common gardens did not provide enough mites for three replicates. Therefore, we only started 37 out of 45 planned population spread assessments with every mesocosm tested at least once. We used similar population spread arenas as the ones in Mortier et al. (2020). A population spread arena consisted of a clean plastic crate (26.5cmx36.5cm) covered in three layers of cotton wool (Rolta®soft) that was kept wet and on which patches of bean leaves (1.5cmx2.5cm) were placed. Bean patches were sequentially connected by a parafilm® bridge (1×8cm) touching both leaves (fig. 2). The remaining leaf’s edges were aligned with paper towel strips (25°C, 16:8h L:D photoperiod).

**Figure 2:**
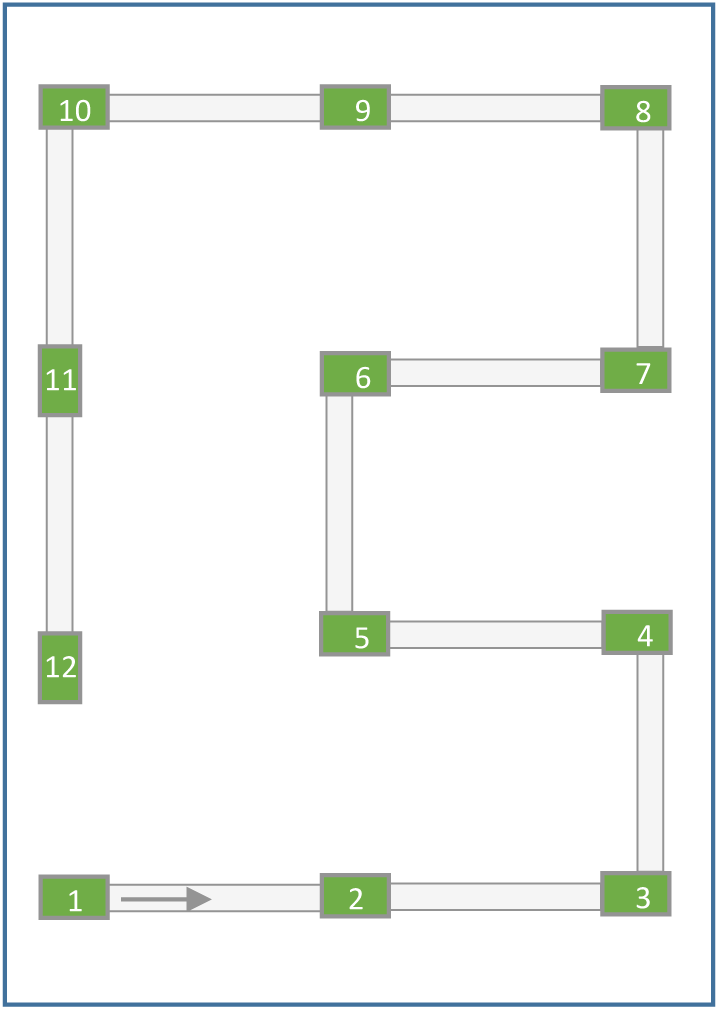
spatial configuration of the population spread arenas. The mites were introduced at patch 1, with possibility to spread beyond the 12^th^ patch

The 40 starting mites were transferred to the first patch with two additional connected empty patches. Every day, if needed, we provided additional empty patches in front as to always ensure two unoccupied patches in front of the furthest occupied patch. This built up the sequence of patches (fig. 2) over the duration of the test. Every two days we replaced the still unoccupied patches at the front to keep the new patches fresh and attractive to arriving mites. This linear patch system snaked through our crates for twelve possible patches. In case of further patches, the first patches and their connection to the next were removed as to provide space for the expanding sequence of patches. We mostly focused on the leading edge of the population distribution and therefore choose to give up trailing patches. In all cases, the removed patch was already withered and did not house any living mites. We kept the population spread arenas at around 25°C for a 16:8h L:D photoperiod. We recorded the furthest occupied patch daily.

### Life history trait tests

We measured dispersal propensity by placing 40 females in their first day of maturity from each common garden, each belonging to a mesocosm, on the first patch in a two-patch dispersal test. The starting bean leaf patch (1.5cmx2.5cm) was connected by a parafilm® bridge (1cmx8cm) to a second patch (25°C, 16:8h L:D photoperiod). This setup tested the number and timing of mites successfully crossing the bridge to the other patch. Every day the destination patch was removed with all successful dispersers of that day to prevent them from going back. A fresh patch is placed to provide an empty destination for the following day. For four days we counted how many individuals still lived and how many successfully dispersed to the second patch to give us a proportion of successfully dispersed individuals. Groups of mites with on average more dispersive traits should have a bigger proportion of the tested mites disperse successfully.

We measured reproductive success by transferring four times a female from each common garden, each belonging to a mesocosm, in the first day of turning adult to a bean leaf patch (1.5cmx2.5cm) on wet cotton wool aligned with paper towel strips (25°C, 16:8h L:D photoperiod). After ten days, we counted the number of adults and deutonymphs (last life stage before adulthood) produced by that female as a measure for her reproductive success.

### Statistical inference

We analyzed all results of our experiment using multilevel modelling and Bayesian estimation methods. The ‘brms’ (Bürkner, 2018) package makes use of ‘Stan’ (Carpenter et al., 2017) as a framework in R in order to estimate posterior parameter distributions using Hamiltonian Monte Carlo (HMC). Replication at multiple levels of the experiment enabled us to estimate the uncertainty on the population spread introduced at the level of the connectedness treatment, the level of the different mesocosms or the replicated assessments of a single mesocosms. This gives us an idea on the relative impact of each level of the experiment on the outcome.

First, we analyzed population spread, the furthest occupied patch, as being dependent on the connectedness treatment the tested mesocosm experienced, time and their interaction with a variable intercept and slope in time for each mesocosm. Second, we modelled population spread the same way but with reproductive success being the focal predictor instead of the historical connectedness treatment. Lastly, we modelled population spread with dispersal propensity as the focal predictor instead of the historical connectedness treatment. In all models we fitted a Gaussian error distribution and used weakly regularizing priors (see supplementary materials).

With the first model, we also calculate the variances accounted for by each predictor or interaction of predictors. In a way we are performing an ANalysis Of VAriance (ANOVA), but in a broad sense. For that, we adapted the method described by Gelman [2007]. The idea is that we can compare the relative impact of predictors and interactions on the outcome by looking at the variation among the predictor’s effect on the outcome, as estimated by the model. We calculated, for each predictor or interaction of predictors, the standard deviation of the estimated marginal effect of that predictor or interaction of predictors on each recorded outcome, so on each data point. We also calculated the estimated residual variation, i.e. the standard deviation in the part of the outcome that is not explained by any predictor or interaction for each data point.

We adapted the method described by (Gelman & Hill, 2007), which calculates the standard deviation of estimated coefficients. Their method has the caveat that estimating the standard deviation among coefficients of an interaction including a continuous variable is affected by the variation in the continuous variables involved. Therefore, this standard deviation is not comparable with standard deviation from main categorical effects. Our method considers the proportional occurrence of each value of a predictor and scales the effect of each predictor and interaction, and the variation therein, to the scale of the outcome.

The data and the script to analyze can be found on https://github.com/fremorti/Evolutionary_history

## Results

When assessing the different sources of variation estimated by the HMC model that predicts population spread from the connectedness treatment, we first notice that the residuals amount to the highest standard deviation (resid, fig. 3). This means that the furthest occupied patch in a test is still varies among observation due to factors not considered. The residual standard deviation is accurately estimated compared the other sources. Furthermore, the time (day) component is an expected source of variation in the spread dynamics. Since mites are obviously introduced in all spread arenas on the starting patch, they could only advance their population edge over time resulting in variation in de furthest occupied patch among different points in time.

**Figure 3:**
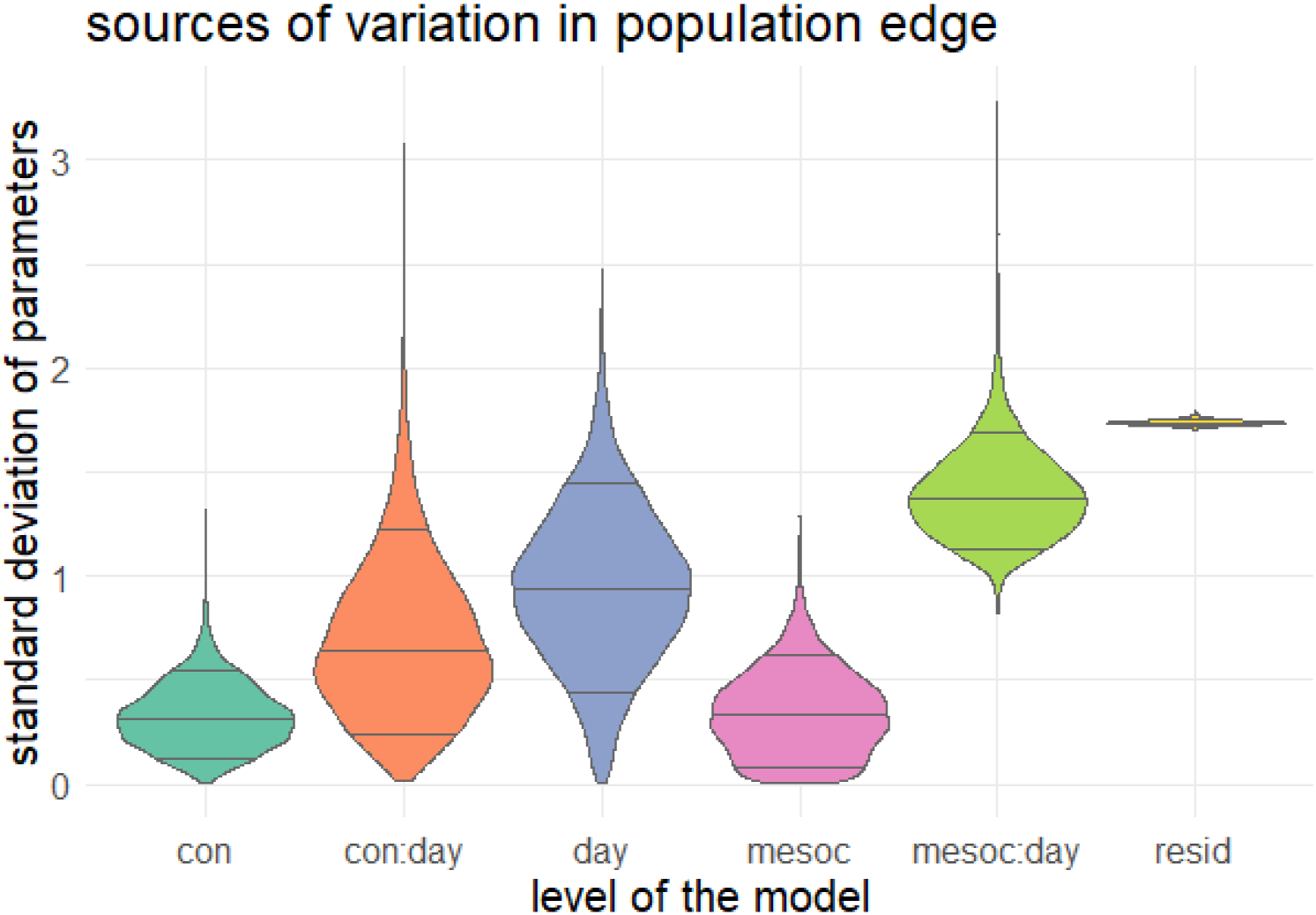
Sources of variation for each parameter, estimated by the HMC model that predicts population spread from the connectedness treatment.

More interestingly, we can compare the variances attributed to the connectedness treatment (con) and to the replicated mesocosms within those treatments (mesoc, fig. 3). The model estimates little variation at the connectedness treatment intercept and mesocosm intercept. Note that all population spread arenas started at the same location, the first patch. In comparison, we see that the estimated interaction with time accounts for a more sizable amount of variance (con:day, mesoc:day; fig.3). We estimate that the slope of the connectedness treatment explains half as much variance in population spread as the slope of the replicated mesocosm itself (fig. 4). The evolutionary history treatment of connectedness is thus accounting for some variation in spread, but differences in the general evolutionary history of the separate mesocosm replicates have a higher impact on spread rate irrespective of their connectedness background.

**Figure 4:**
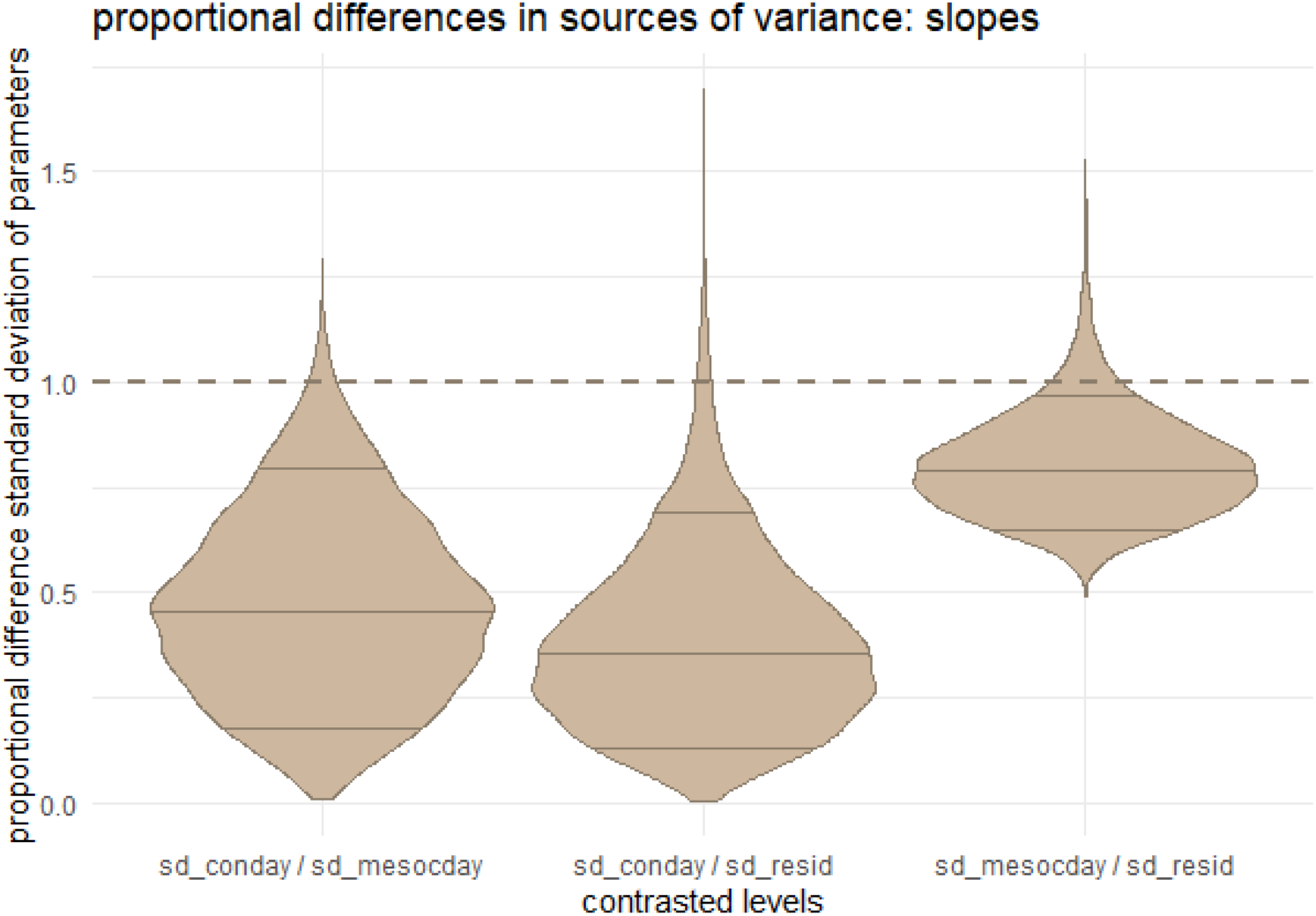
Proportional differences between the estimated variation captured by the connectedness effect on the slope of population spread in time and the mesocosm effect on the slope of population spread in time (left), the proportional difference between the estimated variation captured by the connectedness effect on the slope of population spread in time and the residual variation (middle) and variation captured by the mesocosm effect on the slope of population spread in time and the residual variation (right).

### Fragmentation

The small variation accounted to the fragmentation treatment compared to the mesocosm and residual variation is also nicely illustrated by the unconvincing differences in population spread (fig. 6, left). All treatments show convincingly positive estimated slopes, i.e. population spread rate, but with unconvincing differences between the different connectedness regimes (fig. 5, bottom right).

**Figure 5:**
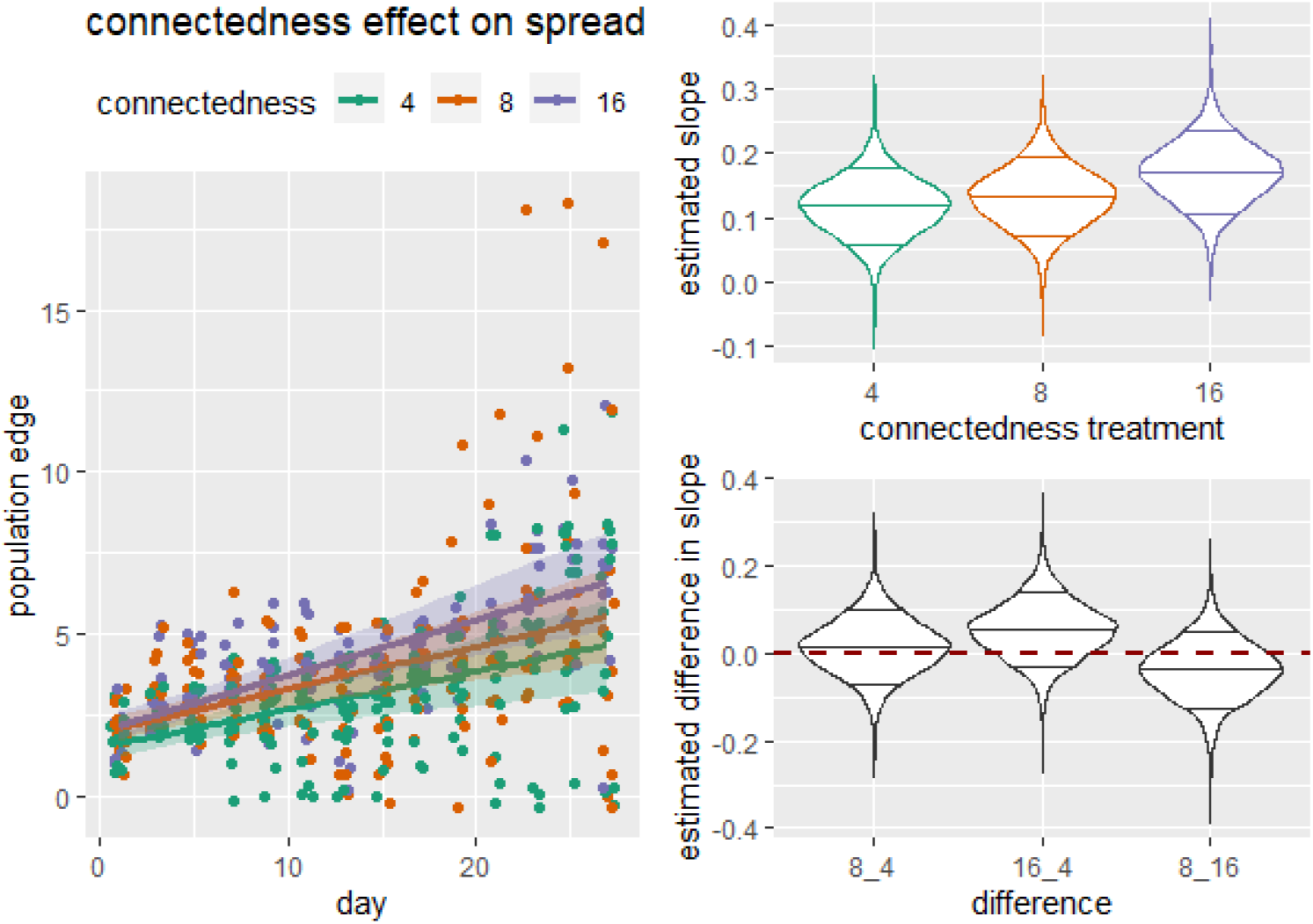
left) The effect on population spread of the connectedness treatment in the evolutionary mesocosms with 4cm (green), 8cm (orange) and 16cm (purple) interpatch distances. Upper right) estimated population spread rate (slope in time) for each connectedness treatment. Lower right) estimated difference in population spread rate between each pair of connectedness treatments.

**Figure 6:**
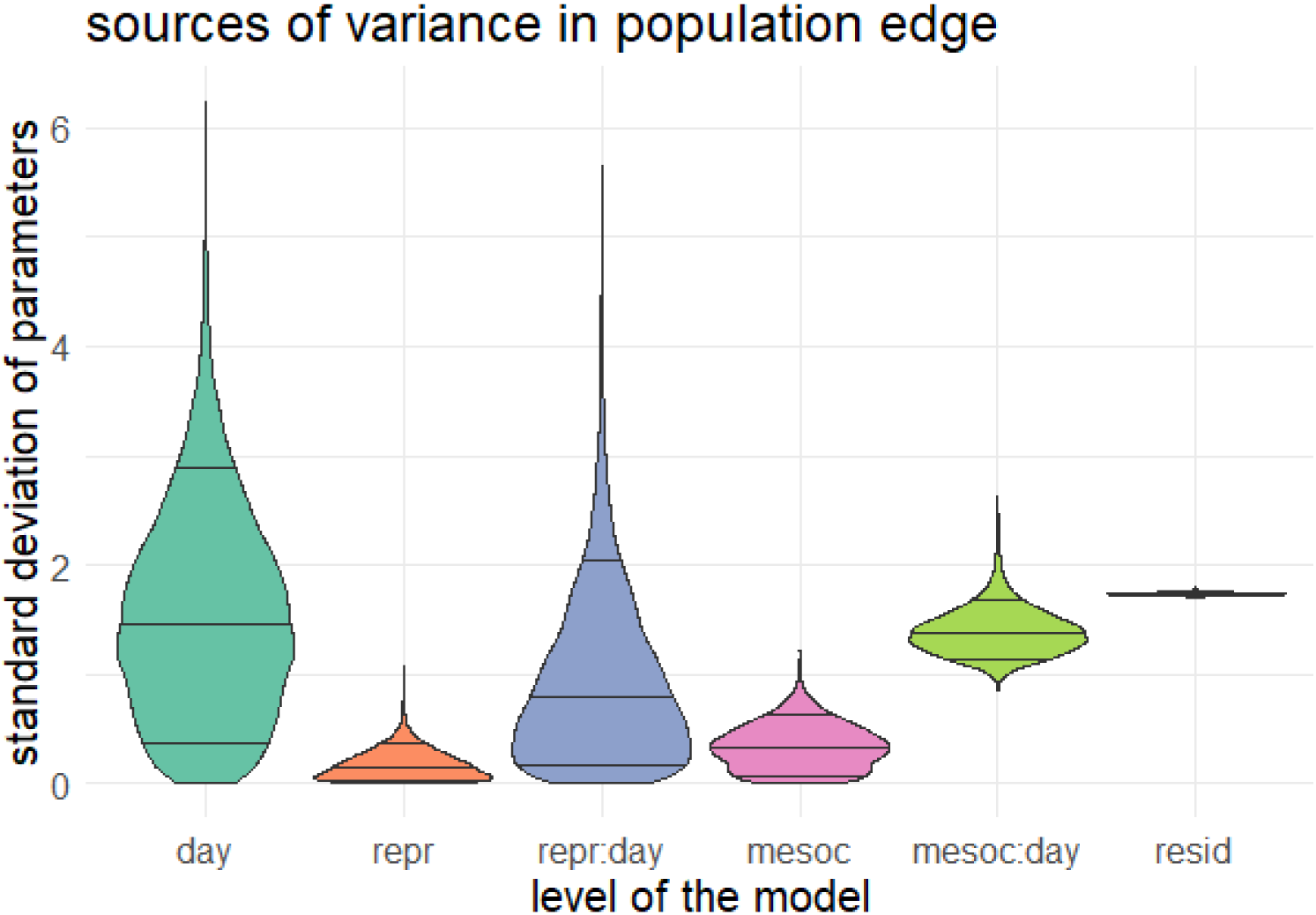
Sources of variation for each parameter, estimated by the HMC model that predicts population spread from the measured reproductive success.

### Role of traits

A portion of the differences in population spread can be attributed to the mesocosm the tested mites originated from. Whether or not this was because of differences in connectedness in the historical environment, it means that the spread rate of a sample of mites resembled that of a different sample of mites from the same mesocosm compared to that of other mesocosms. Therefore, we expect some inherited trait differences that evolved in mesocosms during the evolutionary part of the experiment. We considered two traits that likely affect population spread: reproductive success and dispersal.

### Reproductive success

We estimate a lower amount of variation explained by the interaction of reproductive success and time (repr:day) then for the interaction of the mesocosm and time (mesoc:day). This variation explained is approximately half the residual variation (resid, fig. 6) and is similar to the variation explained by the interaction of connectedness and time (fig. 3). Spread rate, the estimated increase of population edge in time, on average decreases with a higher reproduction but does so unconvincingly (fig. 7).

**Figure 7:**
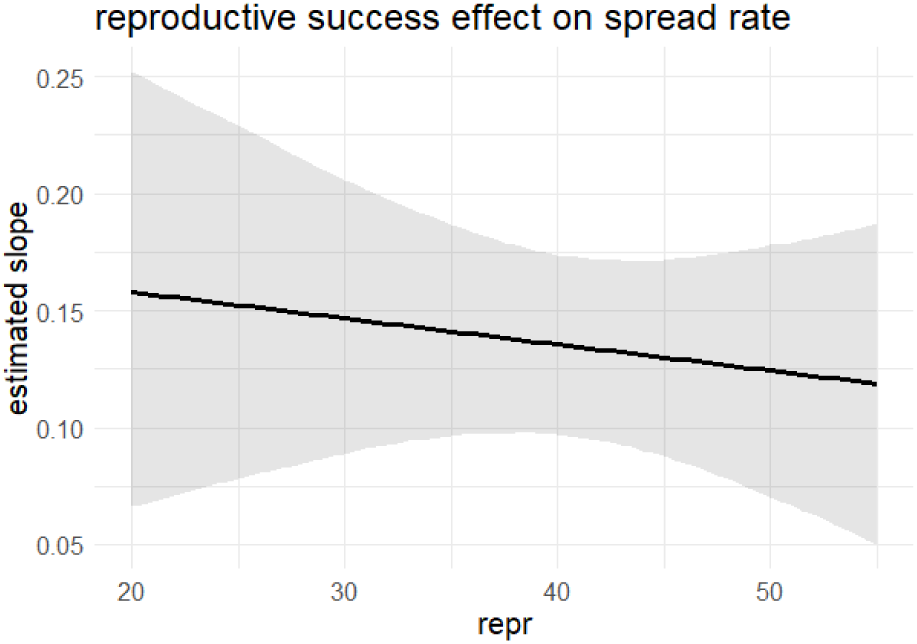
The estimated population spread rate (slope in time) conditional on reproductive success of that population

### Dispersal propensity

We estimate a similar amount of variation explained by the interaction of dispersal propensity and time (disp:day) as by the interaction of the mesocosm and time (mesoc:day). Both variance components are only slightly lower than the residual variation (resid, fig. 8) and relatively higher than the variation explained by the interaction of connectedness and in that model (fig. 3). Paradoxically, spread rate is convincingly lower in populations that evolved a higher dispersal rate (fig. 9).

**Figure 8:**
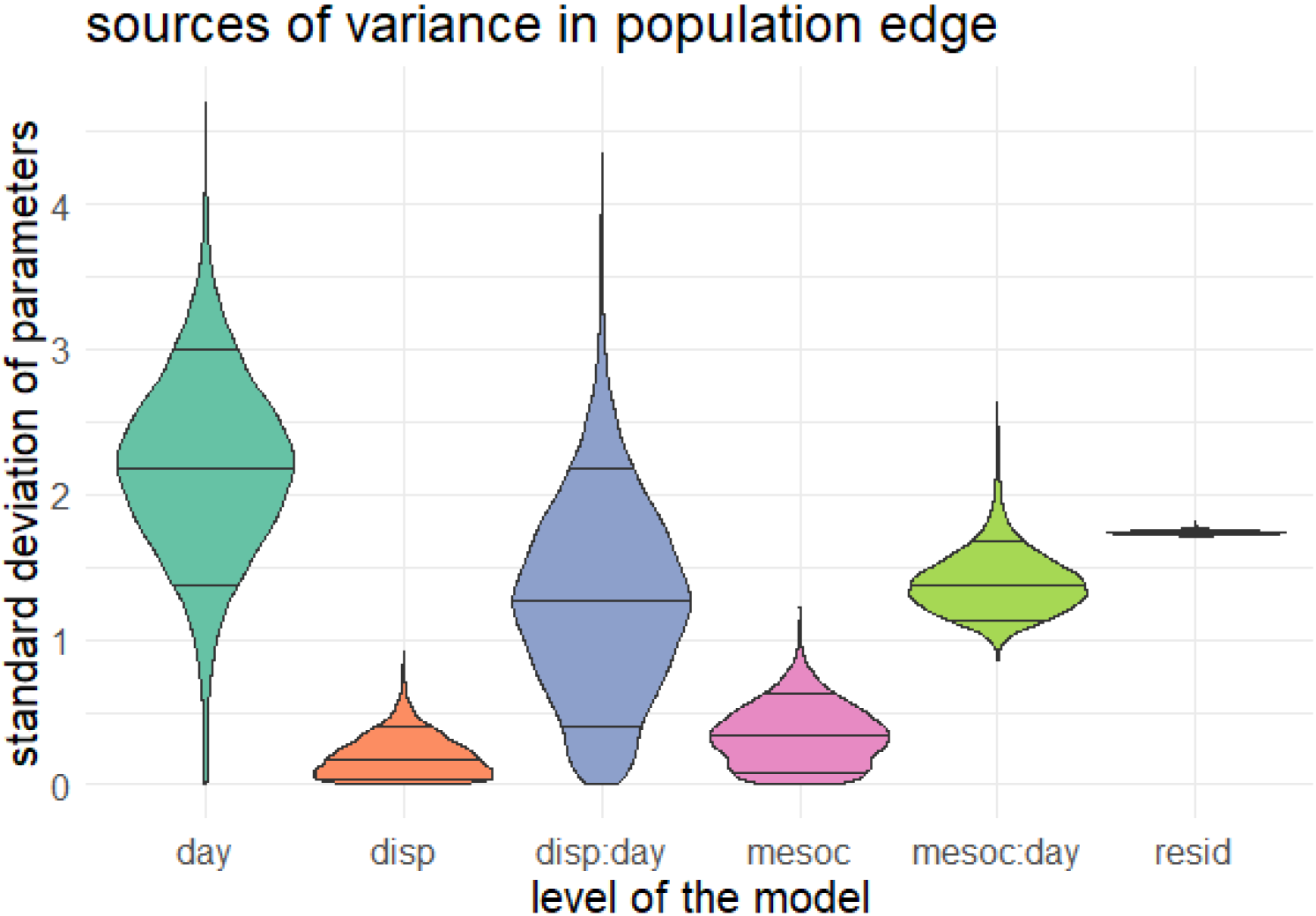
Sources of variation for each parameter, estimated by the HMC model that predicts population spread from the measured dispersal propensity.

**Figure 9:**
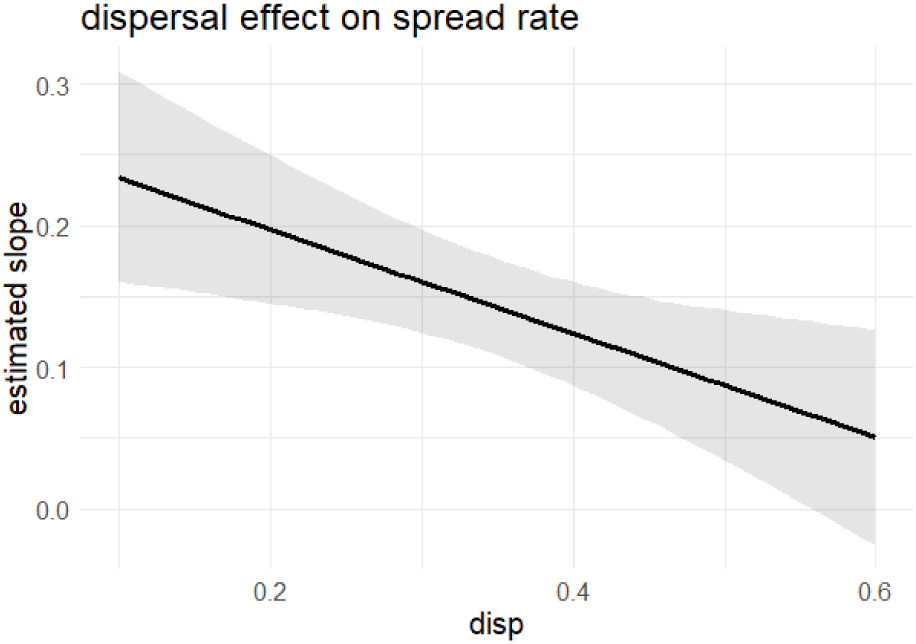
The estimated population spread rate (slope in time) conditional on dispersal propensity of that population

## Discussion

As is often the case in ecological studies, a large component of variation in population spread is left unexplained by our studied predictors. Individual variability and a high level in stochasticity drive individual behavior independent of treatments or other factors on the group level. However, many definable sources of variance contribute considerably to the observed population spread in our experimental population spread. The temporal dimension is here a more trivial source of variation. In time, our mites increased their occupied number of patches when spreading away from the starting patch.

Contrary to our expectations, the evolutionary connectedness treatment encapsulates a rather small amount of variation in population spread dynamics. The mild effect of this deemed relevant evolutionary treatment implies that a population’s ability to outrun environmental change and risk of becoming an invasive species is nevertheless almost impossible to predict from its level of connectedness prior to the population spread. We observe a slight trend of populations from less connected mesocosms to spread faster. This is seemingly at odds with the evolved delayed dispersal at the end of the experimental evolution period (Masier & Bonte, 2019), but we will discuss possible mismatches between dispersal and population spread further below. However, the variation captured by the differences in interpatch distances in the ancestral landscape pales in comparison to the variation captured the variation left unexplained in the analysis.

Interestingly, the experimental mesocosm level encapsulates approximately double the amount of variation compared to the connectedness treatment. We recall that the mesocosm level refers to the replicated mesocosms nested within each connectedness treatment, and each mesocosm in their turn has replicated measurements of population spread. This indicates that populations that experience a similar level of connectedness in their evolutionary history, differ consistently from each other in terms of their potential spread rate. Since all these mesocosms were initialized from the same stock, they must have diverged during the eighteen months of experimental evolution. Since all populations evolved under the same laboratory conditions, with exception of the connectedness treatment, we reason that the relatively large amount of variation attributed to the mesocosm level is predominantly the result of stochastic evolution. This stochastic evolution is as much part of the evolutionary history as the difference in connectedness but is useless when trying to predict future ecological dynamics from it.

While earlier research showed diverging evolution of multiple life history traits in relation to the connectedness background, quite some variation remains within each of these treatments (Masier & Bonte 2020). Hence, evolved traits within each experimental mesocosm might explain variation in the population spread much better. We therefore tested two candidate traits and found dispersal propensity, but not reproductive success, to show a moderately higher predictive power. The direction of the effects was however surprising, as evolved dispersal decreased the rate at which populations spread in time. This counter-intuitive results can only be explained by the presence of trade-offs not tested here. For instance, earlier research using these model organisms found that individuals with a lower tendency to disperse were able to disperse further at the same time (Fronhofer, Stelz, Lutz, Poethke, & Bonte, 2014).

Our study reveals the consistent difficulty to accurately predict the success and extent of population spread (Melbourne & Hastings, 2009). As is often the case in ecological or evolutionary research, the outcome of an experiment or any other repeated observation varies due to stochasticity as a result of sampling or timing of individual events (Cleland, 2001; Pigliucci, 2010). It is the balance between stochastic, chaotic factors and deterministic factors related to the encoding and use of information that determine to what extent we can describe and predict the order in a natural system (O’Connor et al. 2019). Ecological and evolutionary patterns are also hard to predict *a priori* but many times more manageable to explain *a posteriori* when this stochasticity ‘collapses’ into an observation. This ‘asymmetry of overdetermination’ (Cleland, 2001) makes that many patterns of population spread and successful invasions could be explained or rather correlated to features of the organism and environment but that very few generalizations in terms of forecasting can be made (Clark, Lewis, McLachlan, & HilleRisLambers, 2003; Melbourne & Hastings, 2009). We here show that even under standardized laboratory conditions, stochasticity rather than contingency in relation to the environment of origin or expected trait evolution, remains a dominant factor for the eventual outcome of spread dynamics.

Inspired by the predictive power of many physics disciplines and molecular biology, ecologist seek to develop robust forecasting approaches, especially in the field of biodiversity change and invasion biology (Giometto et al., 2014; Melbourne & Hastings, 2009). Our study is another reminder that this will not be easily found as stochasticity and historicity have a big impact on ecological outcome relative to identified tangible drivers of these ecological dynamics (Maris et al., 2018; Pigliucci, 2002). We would like to stress that this incapability of making predictions does not make the field of ecology scientifically any worse at describing reality, the general goal of a science. Hedges (1987) studied replicability, a measure which is thought to be higher in sciences that more successfully describe the world, in the social sciences. Social sciences lend themselves even less to prediction due to the same sources of unpredictability. They nicely revealed that social sciences get on average as consistent results as physics. The difference lies in how variation in results are attributed exclusively to experimental error in physics compared to a myriad of sources of variation in social sciences, usually referred to as the context. Such a context appears to be as important in ecology and evolutionary biology. Instead of trying to achieve generally perfect forecasting, we think it will be more useful to gather insights into the relative magnitude of the sources of variation in ecological and evolutionary dynamics in order to identify in which context determinism dominates and in which contexts forecasting may prove impossible.

## Acknowledgements

We thank Eliane Van der Cruyssen, Kaat Mertens and Noëmie Van den Bon by partaking in the population spread assessments as a part of their bachelor’s dissertation. Additionally, FM thanks the Special Research Fund (BOF) of Ghent University for a PhD scholarship. SM thanks Fonds Wetenschappelijk Onderzoek (FWO) of Flanders for a PhD scholarship. DB, FM and SM are additionally supported by FWO research grant G018017N.

## Data transparency

We provide the data and scripts to analyze at https://github.com/fremorti/Evolutionary_history

## References

Alzate, A., Bisschop, K., Etienne, R. S., & Bonte, D. (2017). Interspecific competition counteracts negative effects of dispersal on adaptation of an arthropod herbivore to a new host. Journal of Evolutionary Biology, 30(11), 1966–1977. https://doi.org/10.1111/jeb.13123

Angert, A. L., Crozier, L. G., Rissler, L. J., Gilman, S. E., Tewksbury, J. J., & Chunco, A. J. (2011). Do species’ traits predict recent shifts at expanding range edges ? Ecology Letters, 14, 677–689. https://doi.org/10.1111/j.1461-0248.2011.01620.x

Bisschop, K., Mortier, F., Etienne, R. S., & Bonte, D. (2019). Transient local adaptation and source– sink dynamics in experimental populations experiencing spatially heterogeneous environments. Proceedings of the Royal Society B: Biological Sciences, 286(1905), 20190738. https://doi.org/10.1098/rspb.2019.0738

Bonte, D., Van Dyck, H., Bullock, J. M., Coulon, A., Delgado, M., Gibbs, M., … Travis, J. M. J. (2012). sts of dispersal. Biological Reviews of the Cambridge Philosophical Society, 87(2), 290–312. https://doi.org/10.1111/j.1469-185X.2011.00201.x

Bürkner, P.-C. (2018). Advanced Bayesian Multilevel Modeling with the R Package brms. The R Journal, 10(1), 395. https://doi.org/10.32614/RJ-2018-017

Burton, O. J., Phillips, B. L., & Travis, J. M. J. (2010). Trade-offs and the evolution of life-histories during range expansion. Ecology Letters, 13, 1210–1220. https://doi.org/10.1111/j.1461-0248.2010.01505.x

Carpenter, B., Gelman, A., Hoffman, M. D., Lee, D., Goodrich, B., Betancourt, M., … Riddell, A. (2017). Stan : A Probabilistic Programming Language. Journal of Statistical Software, 76(1). https://doi.org/10.18637/jss.v076.i01

Cheptou, P. O., Hargreaves, A. L., Bonte, D., & Jacquemyn, H. (2017). Adaptation to fragmentation: Evolutionarydynamics driven by human influences. Philosophical Transactions of the Royal Society B: Biological Sciences, 372(1712). https://doi.org/10.1098/rstb.2016.0037

Clark, J. S., Lewis, M., McLachlan, J. S., & HilleRisLambers, J. (2003). Estimating population spread: What can we forecast and how well? Ecology, 84(8), 1979–1988. https://doi.org/10.1890/01-0618

Cleland, C. E. (2001). Historical science, experimental science, and the scientific method. Geology, 29(11), 987–990. https://doi.org/10.1130/0091-7613(2001)029<0987:HSESAT>2.0.CO;2

De Roissart, A., Wang, S., & Bonte, D. (2015). Spatial and spatiotemporal variation in metapopulation structure affects population dynamics in a passively dispersing arthropod. Journal of Animal Ecology, n/a-n/a. https://doi.org/10.1111/1365-2656.12400

Fisher, R. A. (1937). The wave of advance of advantageous genes. Annals of Eugenics, 7, 355–369.

Fronhofer, E. A., & Altermatt, F. (2015). Eco-evolutionary feedbacks during experimental range expansions. Nature Communications, 6, 6844. https://doi.org/10.1038/ncomms7844

Fronhofer, E. A., Stelz, J. M., Lutz, E., Poethke, H. J., & Bonte, D. (2014). SPATIALLY CORRELATED EXTINCTIONS SELECT FOR LESS EMIGRATION BUT LARGER DISPERSAL DISTANCES IN THE SPIDER MITE TETRANYCHUS URTICAE. Evolution, 68(6), 1838–1844. https://doi.org/10.1111/evo.12339

Gelman, A., & Hill, J. (2007). Data analysis using regression and multilevel/hierarchical models. Cambridge University Press.

Giometto, A., Rinaldo, A., Carrara, F., & Altermatt, F. (2014). Emerging predictable features of replicated biological invasion fronts. 111(1), 297–301. https://doi.org/10.1073/pnas.1321167110

Grbić, M., Van Leeuwen, T., Clark, R. M., Rombauts, S., Rouzé, P., Grbić, V., … Van de Peer, Y. (2011). The genome of Tetranychus urticae reveals herbivorous pest adaptations. (V). https://doi.org/10.1038/nature10640

Hedges, L. V. (1987). How Hard Is Hard Science, How Soft Is Soft Science : The Empirical Cumulativeness of Research. American Psychologist, 42(5), 443–455. https://doi.org/10.1037/0003-066X.42.5.443

Hendry, A. P. (2016). Eco-evolutionary dynamics. Princeton university Press.

Ingvarsson, P. K. (2001). Restoration of genetic variation lost - The genetic rescue hypothesis. Trends in Ecology and Evolution, 16(2), 62–63. https://doi.org/10.1016/S0169-5347(00)02065-6

Jeltsch, F., Bonte, D., Pe’er, G., Reineking, B., Leimgruber, P., Balkenhol, N., … Bauer, S. (2013). Integrating movement ecology with biodiversity research - exploring new avenues to address spatiotemporal biodiversity dynamics. Movement Ecology, 1(1), 6. https://doi.org/10.1186/2051-3933-1-6

Lenormand, T. (2002). Gene flow and the limits to natural selection. Trends in Ecology and Evolution, Vol. 17, pp. 183–189. https://doi.org/10.1016/S0169-5347(02)02497-7

Maris, V., Huneman, P., Coreau, A., Kéfi, S., Pradel, R., & Devictor, V. (2018). Prediction in ecology: promises, obstacles and clarifications. Oikos, 127(2), 171–183. https://doi.org/10.1111/oik.04655

Masier, S., & Bonte, D. (2019). Spatial connectedness imposes local- and metapopulation-level selection on life history through feedbacks on demography. Ecology Letters. https://doi.org/10.1111/ele.13421

Mckinney, M. L., & Lockwood, J. L. (1999). Biotic homogenization : a few winners replacing many losers in the next mass extinction. 5347(Table 1), 450–453.

Melbourne, B. A., & Hastings, A. (2009). Highly variable spread rates in replicated biological invasions: fundamental limits to predictability. Science, 325(September), 1536–1540.

Navajas, M., Perrot-Minnot, M., Lagnel, J., Migeon, A., Bourse, T., & Cornuet, J. M. (2002). Genetic structure of a greenhouse population of the spider mite Tetranychus urticae : spatio-temporal analysis with microsatellite markers. Insect Molecular Biology, 11, 157–165.

O’Connor, M. I., Selig, E. R., Pinsky, M. L., & Altermatt, F. (2012). Toward a conceptual synthesis for climate change responses. Global Ecology and Biogeography, 21(7), 693–703. https://doi.org/10.1111/j.1466-8238.2011.00713.x

Parmesan, C. (2006). Ecological and Evolutionary Responses to Recent Climate Change. Annual Review of Ecology, Evolution, and Systematics, 37(1), 637–669. https://doi.org/10.1146/annurev.ecolsys.37.091305.110100

Pigliucci, M. (2002). Are ecology and evolutionary biology “soft” sciences? Annales Zoologici Fennici, 39(June), 87–98.

Pigliucci, M. (2010). Hard science, soft science. In Nonsense on stilts (pp. 6–23). The University of Chicago Press.

Renault, D., Laparie, M., McCauley, S. J., & Bonte, D. (2018). Environmental Adaptations, Ecological Filtering, and Dispersal Central to Insect Invasions. Annual Review of Entomology, 63(1), 345–368. https://doi.org/10.1146/annurev-ento-020117-043315

Shine, R., Brown, G. P., & Phillips, B. L. (2011). An evolutionary process that assembles phenotypes through space rather than through time. Proceedings of the National Academy of Sciences of the United States of America, 108(14), 5708–5711. https://doi.org/10.1073/pnas.1018989108

Szücs, M., Vahsen, M. L., Melbourne, B. A., Hoover, C., Weiss-Lehman, C., Hufbauer, R. A., & Schoener, T. W. (2017). Rapid adaptive evolution in novel environments acts as an architect of population range expansion. Proceedings of the National Academy of Sciences of the United States of America, 114(51), 13501–13506. https://doi.org/10.1073/pnas.1712934114

Tischendorf, L., & Lenore, G. (2001). On the Use of Connectivity Measures in Spatial Ecology. A Reply. Oikos, 95(1), 152–155.

Van Petegem, K., Moerman, F., Dahirel, M., Fronhofer, E. A., Vandegehuchte, M. L., Van Leeuwen, T., … Bonte, D. (2018). Kin competition accelerates experimental range expansion in an arthropod herbivore. Ecology Letters, 21, 225–234. https://doi.org/10.1111/ele.12887

