## supplementary materials for "Evolutionary history determines population spread rate in a stochastic, rather than in a deterministic way"

### Appendix: Statistical model description and estimates

All models were run in R using brms and Hamiltonian Monte Carlo with two chains with each 5000 iteration from which 2000 were warmup

#### Model 1: Population spread depending on connectedness treatment

a) Model representation

edge ~ 0 + Intercept + treatment*day + (1 + day|mesoc)

prior = c(prior(normal(0,4), class = b),

prior(cauchy(0,2), class = sd),

prior(lkj(2), class = cor)

With *edge* the furthest occupied patch in a population spread test of a sample originating from an experimental mesocosm (*mesoc*) of a certain connectedness *treatment* spreading recorded at a certain point in time (in number of *day*s since the start). We modelled edge with a normally distributed error distribution and estimated it depending on *treatment*, time (*day*) and their interaction. We also modelled a varying intercept and slope in time for each tested mesocosm (*mesoc*). We used priors that are weakly regularizing. Of note, the used LKJ distribution is a Lewandowski-Kurowicka-Joe distribution that is regularly used a prior for a correlation matrix.

b) Model parameter estimates

| | mean | se_mean | sd | 2.5% | 97.5% | n_eff | Rhat |
|---|---|---|---|---|---|---|---|
| b_Int | 1.49 | 0.01 | 0.31 | 0.88 | 2.12 | 3492 | 1 |
| b_treatment8 | 0.48 | 0.01 | 0.45 | -0.39 | 1.34 | 4291 | 1 |
| b_treatment16 | 0.51 | 0.01 | 0.49 | -0.48 | 1.42 | 3963 | 1 |
| b_day | 0.12 | 0.00 | 0.05 | 0.02 | 0.21 | 2173 | 1 |
| b_treatment8:day | 0.02 | 0.00 | 0.07 | -0.12 | 0.15 | 2434 | 1 |
| b_treatment16:day | 0.05 | 0.00 | 0.07 | -0.08 | 0.19 | 2566 | 1 |
| sd_mesoc__Int | 0.38 | 0.01 | 0.24 | 0.02 | 0.94 | 2151 | 1 |
| sd_mesoc__day | 0.10 | 0.00 | 0.02 | 0.06 | 0.16 | 1947 | 1 |
| cor_mesoc_Int_day | -0.41 | 0.01 | 0.38 | -0.91 | 0.52 | 931 | 1 |
| sigma | 1.74 | 0.00 | 0.05 | 1.64 | 1.85 | 10190 | 1 |
| r_mesoc[16.1,Int] | -0.02 | 0.00 | 0.35 | -0.79 | 0.67 | 6434 | 1 |
| r_mesoc[16.2,Int] | 0.30 | 0.01 | 0.40 | -0.28 | 1.29 | 3808 | 1 |
| r_mesoc[16.3,Int] | -0.03 | 0.00 | 0.36 | -0.78 | 0.74 | 5675 | 1 |
| r_mesoc[16.4,Int] | -0.12 | 0.01 | 0.39 | -1.05 | 0.56 | 5146 | 1 |

| | | | | | | | |
|---|---|---|---|---|---|---|---|
| r_mesoc[16.5,Int] | -0.12 | 0.00 | 0.37 | -0.95 | 0.57 | 6040 | 1 |
| r_mesoc[4.1,Int] | -0.19 | 0.01 | 0.35 | -1.03 | 0.42 | 3459 | 1 |
| r_mesoc[4.2,Int] | -0.20 | 0.01 | 0.36 | -1.07 | 0.40 | 3686 | 1 |
| r_mesoc[4.3,Int] | -0.02 | 0.01 | 0.34 | -0.72 | 0.73 | 4025 | 1 |
| r_mesoc[4.4,Int] | 0.08 | 0.00 | 0.33 | -0.58 | 0.79 | 4367 | 1 |
| r_mesoc[4.5,Int] | 0.32 | 0.01 | 0.42 | -0.31 | 1.33 | 3326 | 1 |
| r_mesoc[8.1,Int] | -0.40 | 0.01 | 0.44 | -1.38 | 0.25 | 2575 | 1 |
| r_mesoc[8.2,Int] | 0.22 | 0.01 | 0.41 | -0.44 | 1.23 | 4728 | 1 |
| r_mesoc[8.3,Int] | 0.03 | 0.00 | 0.33 | -0.65 | 0.72 | 6780 | 1 |
| r_mesoc[8.4,Int] | 0.18 | 0.00 | 0.34 | -0.42 | 0.99 | 5345 | 1 |
| r_mesoc[8.5,Int] | -0.02 | 0.00 | 0.33 | -0.74 | 0.64 | 6535 | 1 |
| r_mesoc[16.1,day] | -0.05 | 0.00 | 0.05 | -0.15 | 0.05 | 4165 | 1 |
| r_mesoc[16.2,day] | -0.03 | 0.00 | 0.05 | -0.13 | 0.07 | 3799 | 1 |
| r_mesoc[16.3,day] | 0.07 | 0.00 | 0.05 | -0.03 | 0.18 | 4427 | 1 |
| r_mesoc[16.4,day] | -0.03 | 0.00 | 0.05 | -0.14 | 0.07 | 4278 | 1 |
| r_mesoc[16.5,day] | 0.04 | 0.00 | 0.05 | -0.07 | 0.15 | 4464 | 1 |
| r_mesoc[4.1,day] | 0.08 | 0.00 | 0.05 | -0.02 | 0.18 | 2449 | 1 |
| r_mesoc[4.2,day] | 0.01 | 0.00 | 0.05 | -0.09 | 0.11 | 2439 | 1 |
| r_mesoc[4.3,day] | 0.09 | 0.00 | 0.05 | -0.01 | 0.19 | 2473 | 1 |
| r_mesoc[4.4,day] | -0.07 | 0.00 | 0.05 | -0.17 | 0.03 | 2473 | 1 |
| r_mesoc[4.5,day] | -0.10 | 0.00 | 0.05 | -0.21 | -0.01 | 2650 | 1 |
| r_mesoc[8.1,day] | 0.17 | 0.00 | 0.05 | 0.08 | 0.28 | 3401 | 1 |
| r_mesoc[8.2,day] | -0.07 | 0.00 | 0.05 | -0.19 | 0.03 | 3882 | 1 |
| r_mesoc[8.3,day] | -0.03 | 0.00 | 0.05 | -0.13 | 0.06 | 4027 | 1 |
| r_mesoc[8.4,day] | -0.03 | 0.00 | 0.05 | -0.13 | 0.07 | 4047 | 1 |
| r_mesoc[8.5,day] | -0.04 | 0.00 | 0.05 | -0.13 | 0.06 | 4064 | 1 |

### Model 2: Population spread variance

a) Model representation

edge ~ edge ~ 0 + Intercept + repr*day + (1+day|mesoc)

prior = c(prior(normal(0,4), class = b),

prior(cauchy(0,2), class = sd),

prior(lkj(2), class = cor)

With *edge* the furthest occupied patch in a population spread test of a sample originating from an experimental mesocosm (*mesoc*) that exhibited a certain reproductive success (*repr*) recorded at a certain point in time (in number of *day*s since the start). We modelled edge with a normally distributed error distribution and estimated it depending on *repr*oductive success, time (*day*) and their interaction. We also modelled a varying intercept and slope in time for each tested mesocosm (*mesoc*). We used priors that are weakly regularizing. Of note, the used LKJ distribution is a Lewandowski-Kurowicka-Joe distribution that is regularly used a prior for a correlation matrix.

b) Model parameter estimates

| | mean | se_mean | sd | 2.5% | 97.5% | n_eff | Rhat |
|---|---|---|---|---|---|---|---|
| b_Int | 1.39 | 0.02 | 0.86 | -0.37 | 3.06 | 3023 | 1 |
| b_repr | 0.01 | 0.00 | 0.02 | -0.03 | 0.05 | 3067 | 1 |
| b_day | 0.19 | 0.00 | 0.13 | -0.08 | 0.46 | 2304 | 1 |
| b_repr:day | 0.00 | 0.00 | 0.00 | -0.01 | 0.00 | 2229 | 1 |
| sd_mesoc__Int | 0.39 | 0.01 | 0.27 | 0.02 | 0.99 | 2015 | 1 |
| sd_mesoc__day | 0.10 | 0.00 | 0.03 | 0.06 | 0.17 | 1722 | 1 |
| cor_mesoc_Int_day | -0.20 | 0.01 | 0.40 | -0.83 | 0.65 | 832 | 1 |
| sigma | 1.81 | 0.00 | 0.06 | 1.70 | 1.93 | 6092 | 1 |
| r_mesoc[16.1,Int] | -0.03 | 0.00 | 0.32 | -0.70 | 0.64 | 6130 | 1 |
| r_mesoc[16.2,Int] | 0.32 | 0.01 | 0.41 | -0.24 | 1.33 | 3071 | 1 |
| r_mesoc[16.3,Int] | 0.05 | 0.01 | 0.37 | -0.66 | 0.92 | 4343 | 1 |
| r_mesoc[16.4,Int] | -0.18 | 0.01 | 0.41 | -1.19 | 0.52 | 4290 | 1 |
| r_mesoc[4.2,Int] | -0.25 | 0.01 | 0.40 | -1.22 | 0.38 | 3609 | 1 |
| r_mesoc[4.3,Int] | -0.07 | 0.00 | 0.33 | -0.85 | 0.56 | 4510 | 1 |
| r_mesoc[4.4,Int] | -0.06 | 0.01 | 0.34 | -0.85 | 0.63 | 4295 | 1 |
| r_mesoc[4.5,Int] | 0.16 | 0.01 | 0.39 | -0.53 | 1.12 | 3056 | 1 |
| r_mesoc[8.1,Int] | -0.25 | 0.01 | 0.40 | -1.17 | 0.41 | 2592 | 1 |
| r_mesoc[8.2,Int] | 0.17 | 0.01 | 0.42 | -0.55 | 1.19 | 4230 | 1 |
| r_mesoc[8.3,Int] | 0.00 | 0.00 | 0.33 | -0.74 | 0.71 | 5642 | 1 |
| r_mesoc[8.4,Int] | 0.27 | 0.01 | 0.43 | -0.35 | 1.35 | 2636 | 1 |
| r_mesoc[8.5,Int] | -0.08 | 0.01 | 0.35 | -0.86 | 0.62 | 4466 | 1 |
| r_mesoc[16.1,day] | 0.00 | 0.00 | 0.04 | -0.08 | 0.07 | 4052 | 1 |
| r_mesoc[16.2,day] | 0.02 | 0.00 | 0.04 | -0.06 | 0.09 | 3312 | 1 |
| r_mesoc[16.3,day] | 0.11 | 0.00 | 0.04 | 0.03 | 0.20 | 4053 | 1 |
| r_mesoc[16.4,day] | 0.02 | 0.00 | 0.05 | -0.08 | 0.12 | 3815 | 1 |
| r_mesoc[4.2,day] | -0.03 | 0.00 | 0.06 | -0.15 | 0.09 | 2723 | 1 |
| r_mesoc[4.3,day] | 0.06 | 0.00 | 0.04 | -0.02 | 0.14 | 3436 | 1 |
| r_mesoc[4.4,day] | -0.10 | 0.00 | 0.04 | -0.19 | -0.02 | 3199 | 1 |
| r_mesoc[4.5,day] | -0.12 | 0.00 | 0.04 | -0.22 | -0.04 | 3489 | 1 |
| r_mesoc[8.1,day] | 0.17 | 0.00 | 0.04 | 0.10 | 0.26 | 3422 | 1 |
| r_mesoc[8.2,day] | -0.05 | 0.00 | 0.06 | -0.17 | 0.05 | 3022 | 1 |
| r_mesoc[8.3,day] | -0.02 | 0.00 | 0.05 | -0.11 | 0.07 | 3135 | 1 |
| r_mesoc[8.4,day] | -0.04 | 0.00 | 0.07 | -0.19 | 0.10 | 2494 | 1 |
| r_mesoc[8.5,day] | -0.01 | 0.00 | 0.06 | -0.13 | 0.10 | 2726 | 1 |

### Model 3: Total population size

a) Model representation

edge ~ edge ~ 0 + Intercept + disp*day + (1+day|mesoc)

prior = c(prior(normal(0,4), class = b),

prior(cauchy(0,2), class = sd),

prior(lkj(2), class = cor)

With edge the furthest occupied patch in a population spread test of a sample originating from an experimental mesocosm (*mesoc*) that exhibited a certain dispersal propensity (*disp*) recorded at a certain point in time (in number of *day*s since the start). We modelled edge with a normally distributed error distribution and estimated it depending on *disp*ersal propensity, time (day) and their interaction. We also modelled a varying intercept and slope in time for each tested mesocosm (*mesoc*). We used priors that are weakly regularizing. Of note, the used LKJ distribution is a Lewandowski-Kurowicka-Joe distribution that is regularly used a prior for a correlation matrix.

b) Model estimates

| | mean | se_mean | sd | 2.5% | 97.5% | n_eff | Rhat |
|---|---|---|---|---|---|---|---|
| b_Int | 1.44 | 0.01 | 0.59 | 0.28 | 2.60 | 3348 | 1 |
| b_disp | 1.14 | 0.03 | 1.58 | -1.98 | 4.29 | 3301 | 1 |
| b_day | 0.27 | 0.00 | 0.08 | 0.12 | 0.42 | 2472 | 1 |
| b_disp:day | -0.36 | 0.00 | 0.20 | -0.78 | 0.05 | 2471 | 1 |
| sd_mesoc_Int | 0.42 | 0.01 | 0.27 | 0.02 | 1.01 | 2015 | 1 |
| sd_mesoc_day | 0.08 | 0.00 | 0.02 | 0.05 | 0.14 | 1674 | 1 |
| cor_mesoc_Int_day | -0.27 | 0.01 | 0.39 | -0.85 | 0.64 | 908 | 1 |
| sigma | 1.79 | 0.00 | 0.06 | 1.67 | 1.91 | 6491 | 1 |
| r_mesoc[16.1,Int] | -0.06 | 0.00 | 0.34 | -0.80 | 0.62 | 6241 | 1 |
| r_mesoc[16.2,Int] | 0.40 | 0.01 | 0.45 | -0.18 | 1.46 | 3128 | 1 |
| r_mesoc[16.3,Int] | 0.10 | 0.01 | 0.40 | -0.64 | 1.02 | 4696 | 1 |
| r_mesoc[16.4,Int] | -0.16 | 0.01 | 0.41 | -1.13 | 0.53 | 5122 | 1 |
| r_mesoc[4.1,Int] | -0.30 | 0.01 | 0.41 | -1.24 | 0.34 | 3206 | 1 |
| r_mesoc[4.2,Int] | -0.24 | 0.01 | 0.43 | -1.30 | 0.46 | 3261 | 1 |
| r_mesoc[4.3,Int] | 0.02 | 0.01 | 0.36 | -0.75 | 0.81 | 5038 | 1 |
| r_mesoc[4.5,Int] | 0.13 | 0.01 | 0.41 | -0.66 | 1.10 | 4222 | 1 |
| r_mesoc[8.1,Int] | -0.28 | 0.01 | 0.43 | -1.25 | 0.46 | 2233 | 1 |
| r_mesoc[8.2,Int] | 0.21 | 0.01 | 0.43 | -0.50 | 1.30 | 4049 | 1 |
| r_mesoc[8.3,Int] | 0.05 | 0.00 | 0.33 | -0.63 | 0.78 | 6463 | 1 |
| r_mesoc[8.4,Int] | 0.20 | 0.01 | 0.36 | -0.40 | 1.09 | 4528 | 1 |
| r_mesoc[8.5,Int] | -0.05 | 0.00 | 0.34 | -0.80 | 0.61 | 6011 | 1 |
| r_mesoc[16.1,day] | 0.01 | 0.00 | 0.04 | -0.07 | 0.08 | 4392 | 1 |
| r_mesoc[16.2,day] | -0.03 | 0.00 | 0.04 | -0.11 | 0.05 | 3626 | 1 |
| r_mesoc[16.3,day] | 0.05 | 0.00 | 0.05 | -0.04 | 0.15 | 3116 | 1 |
| r_mesoc[16.4,day] | 0.00 | 0.00 | 0.04 | -0.08 | 0.08 | 5085 | 1 |
| r_mesoc[4.1,day] | 0.07 | 0.00 | 0.04 | -0.01 | 0.16 | 3139 | 1 |
| r_mesoc[4.2,day] | -0.08 | 0.00 | 0.05 | -0.19 | 0.02 | 3052 | 1 |
| r_mesoc[4.3,day] | -0.01 | 0.00 | 0.05 | -0.10 | 0.08 | 3115 | 1 |
| r_mesoc[4.5,day] | -0.07 | 0.00 | 0.06 | -0.19 | 0.04 | 3233 | 1 |
| r_mesoc[8.1,day] | 0.16 | 0.00 | 0.04 | 0.10 | 0.24 | 2822 | 1 |
| r_mesoc[8.2,day] | -0.05 | 0.00 | 0.05 | -0.15 | 0.04 | 4371 | 1 |
| r_mesoc[8.3,day] | -0.04 | 0.00 | 0.03 | -0.10 | 0.03 | 5204 | 1 |
| r_mesoc[8.4,day] | -0.01 | 0.00 | 0.04 | -0.09 | 0.06 | 4290 | 1 |
| r_mesoc[8.5,day] | -0.01 | 0.00 | 0.04 | -0.08 | 0.07 | 4271 | 1 |
